# FABP7 Binds to Fatty Acid Micelles: Implications for Lipid Transport

**DOI:** 10.1101/2021.10.22.465361

**Authors:** Stefan Lenz, Iulia Bodnariuc, Margaret Renaud-Young, Tanille M. Shandro, Justin L. MacCallum

**Affiliations:** University of Calgary

**Author notes:** equal first author (Stefan Lenz, Iulia Bodnariuc, Margaret Renaud-Young).

## Abstract

The transport of hydrophobic molecules, including long-chain fatty acids, within cells is highly dynamic. Hydrophobic molecules are unable to freely diffuse through the aqueous cytoplasm without a transporter. Fatty acid binding proteins (FABP) transport these molecules to different cellular compartments. As part of their transport, FABPs often associate with cell membranes to acquire and deliver their bound cargo. Understanding the nature of this transport is becoming increasingly important because lipid signaling functions are associated with metabolic pathways impacting disease pathologies such as carcinomas, autism and schizophrenia. Herein, we focus on Brain fatty acid binding protein (FABP7), which demonstrates localization to the cytoplasm and nucleus, influencing transcription and fatty acid metabolism. We use a combined biophysical approach to elucidate the interaction between FABP7 and model membranes. Specifically, we use microscale thermophoresis to show that FABP7 can bind oleic acid (OA) and docosahexaenoic acid (DHA) micelles, while differential scanning fluorimetry experiments show binding lowers the melting temperature of FABP7. Structural data from NMR and multiscale molecular dynamics simulations reveals that the interaction between FABP7 and micelles is through FABP7’s portal region residues. Our simulations also capture binding events where fatty acids dissociate from the model membrane and bind to FABP7. Overall, our data reveals a novel interaction between FABP7 and OA or DHA micelles and provides key structural insight into the transport of hydrophobic molecules.

**Significance:** This study examines how FABP7 binds to fatty acids at low and high fatty acid concentrations. Our binding assays, including microscale thermophoresis (MST) and Nile red fluorescence establish that FABP7 binds to both free fatty acids in solution and fatty acid micelles. NMR and computational experiments show that FABP7 specifically interacts with micelles through the portal region of the protein, thereby mediating ligand transfer into the binding cavity.

## Introduction

Fatty acid binding proteins (FABPs) are 14-16 kDa cyto- and nucleosolic protein transporters of hydrophobic molecules (1). This transport intersects with many cellular functions, including cell mobility, inflammation, energy storage, and membrane synthesis (2). Nine human isoforms of FABPs have been identified thus far with tissue-specific expression and intercellular transportation.

Depending on their cargo, FABPs can be localized to many sub-cellular organelles, including, for example, the nucleus (3), golgi apparatus (4), peroxisomes (5), and the endoplasmic reticulum (6). Among the nine FABP isoforms is brain FABP (FABP7), which is primarily expressed in brain, glial, retina, and mammary gland cells (7). FABP7 has a high affinity for poly unsaturated long chain fatty acids. In particular, FABP7 has a high affinity for docosahexaenoic acid (22:6, DHA), which leads to increased localization of the protein to the nucleus where it can affect gene expression by activating peroxisome proliferator-activated receptors (PPARs) (8). The role of FABP7 is particularly important during the development of the nervous system where high expression levels of FABP7 are associated with increased cell mobility (9). However, higher expression levels of FABP7 is associated with poor prognosis in patients afflicted with melanoma (10) or carcinoma (11).

Despite modest primary sequence similarity (28 to 70%), FABPs share a common tertiary structure with ten anti-parallel β-strands forming a β barrel and two α-helices (Fig. 1) (12). Within the β barrel is a ∼500 Å cavity where FABPs bind to hydrophobic cargo. Among the structurally diverse array of ligands that FABPs can bind is fatty acids. For FABP7, the carboxylate group of bound fatty acids coordinates to the sidechains of R126 and Y128 (13). Ligand entry and exit is proposed to be mediated by a portal region composed of the α-helices and βCD and βEF loops. In addition to binding molecules within the β barrel, FABPs bind to membranes and membrane mimetics, including micelles (14) and nanodiscs (15). In general, the interaction between FABPs and micelles or micelle mimetics falls under either collisional or diffusional mechanisms (16, 17). Collisional transfer involves association of the FABP with the membrane and direct transfer, while diffusional entails solvent-mediated transfer and no direct interaction between the protein and membrane. Several FABPs, including FABP7, with positively charged alpha helices utilize a collisional mechanism (18–21), while liver FABP (FABP1) uses a diffusional mechanism (22). However, the precise way ligands enter the FABP binding pocket is not known.

**Figure 1.**
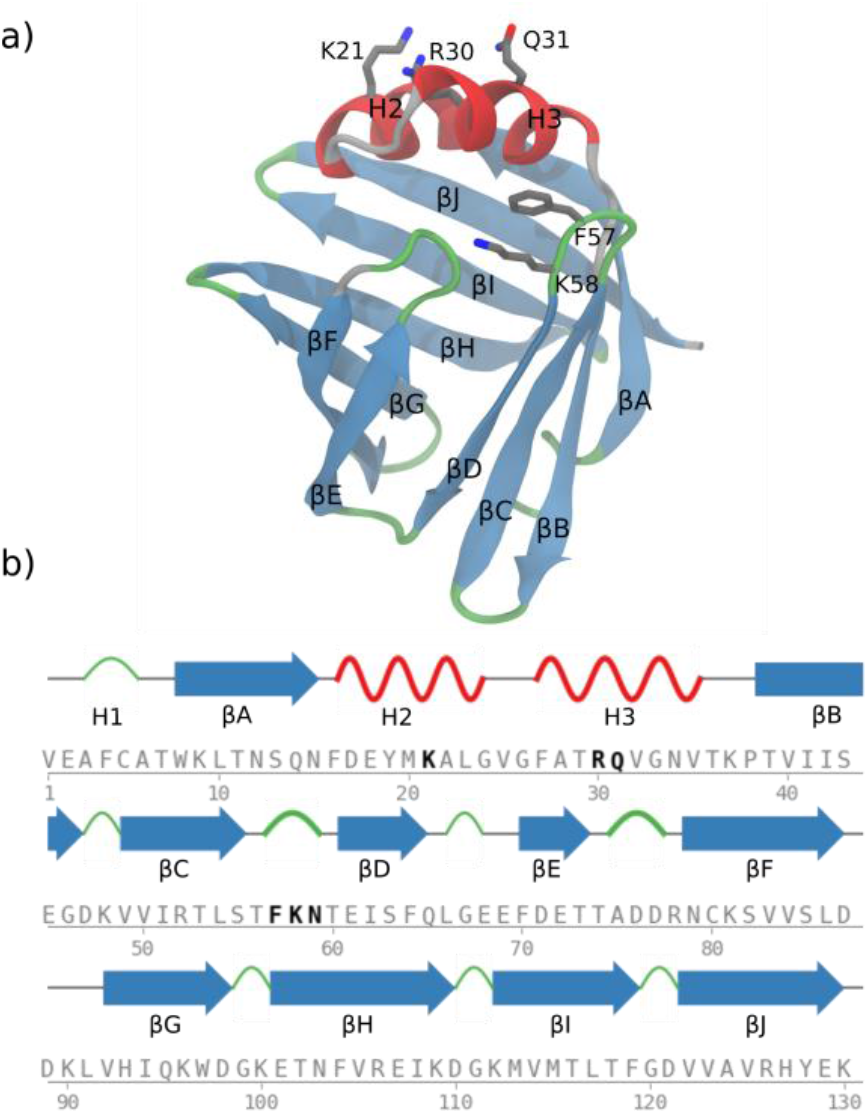
FABP7 structure and sequence with key membrane- and micelle-binding residues highlighted. a) Structure and secondary structure numbering scheme used in the current study. b) FABP7 sequence with secondary structure elements labelled.

Here we highlight a second binding interaction observed between FABP7 and OA or DHA that can only be observed at high ligand concentrations. Within this framework, we see unique changes to the portal region of the protein that are not seen when it interacts with single ligands. We suggest that this mimics FABP7 interactions with membranes and provides insight into the ligand binding/exchange behavior of the protein.

We sought to understand how FABP7 interacts with model membranes and how that influences ligand binding and egress. We use fluorescence intensity change and MST assays to characterize the strength of the binding interaction between FABP7 and micelles. To understand the structural basis for the binding interaction, we first use NMR to examine the chemical shifts differences that manifest in the presence of micelles. Subsequently, we use a multi-scale modelling molecular dynamics approach to examine the specific residues that interact with micelles. We also model the corresponding interaction between FABP7 and zwitterionic or anionic membranes. Our findings are:

- Binding assays reveal a second binding event occurs at ligand concentrations where micelles form (i.e. the critical micelle concentration). Binding models indicate that this event occurs at a greater than 1:1 ligand to protein ratio supporting binding between micelles and FABP7.
- Computer simulations predict a specific binding interaction between FABP7 and micelles involving several residues in the portal region. This interaction also alters the structural dynamics of the protein.
- NMR experiments at different ligand concentrations show that residues in the portal region have diverging behavior above the concentration that single molecule pocket binding occurs. This is indicative of complex equilibria involving changes to protein conformations coupled with ligand and micelle binding.
- Our simulations also predict that FABP7 interacts with membranes in a similar manner to how it interacts with micelles. Interestingly, the binding interaction is stronger with more anionic membranes.
- During our simulations we observe single fatty acids transition from the micelle to FABP7 binding pocket and back. These single molecule binding and unbinding events are mediated by residues in H3 and βCD.

## Results and Discussion

### MST experiments reveal two binding events between FABP7 and unsaturated fatty acids

Microscale thermophoresis (MST) was used to determine the binding affinity of fluorescently-labelled FABP7 (labelled with RED-tris-NHS dye) to the fatty acids using *thermophoresis* – the movement of fluorescently labelled protein in a temperature gradient – and *initial fluorescence* – the change in fluorescence that occurs without a temperature change. We observe no change in fluorescence from a control experiment with the NHS-REDtris fluorophore alone against a dilution series of OA and DHA (Fig. S1).

Our MST binding assays performed at 37 °C reveal a binding event from changes to *thermophoresis* at ∼0.5 μM between FABP7 and each fatty acid: oleic acid (OA), DHA, and stearic acid (SA; Figs. 2 and S2). This corresponds to a single fatty acid molecule entering the binding cavity of FABP7. These binding affinities are generally within an order of magnitude of literature values (23).

**Figure 2.**
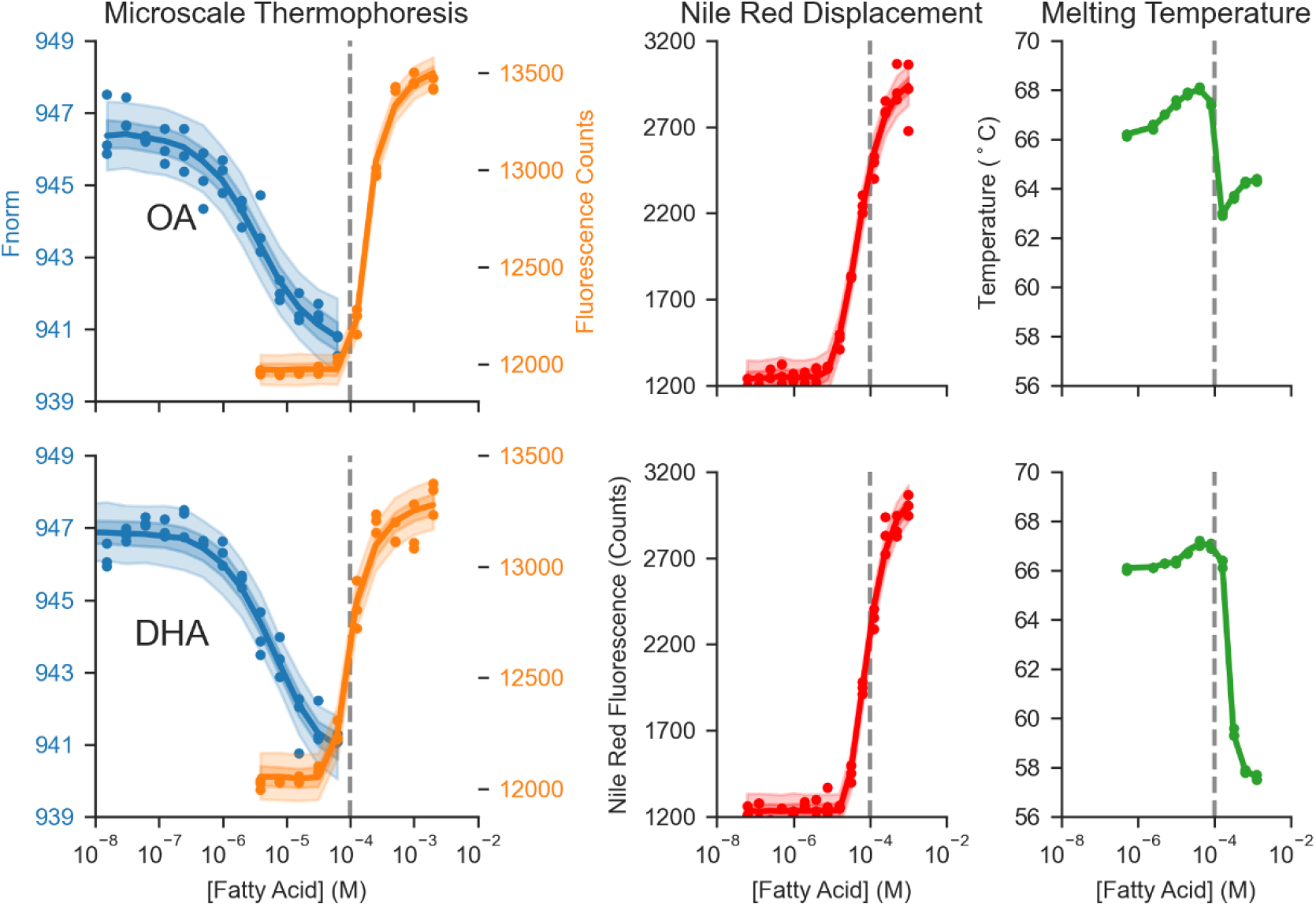
Dilution series experiments probing OA and DHA binding to FABP7 suggests that the protein binds single fatty acids in its pocket and binds to micelles as they form. Thermophoresis (left, blue) and initial fluorescence (left, orange) data for FABP7 binding to OA (top) and DHA (bottom) reveal two binding events. Nile red assay (middle, red) determination of the critical micelle concentration reveals that the second binding events occurs at roughly the same concentration that micelles form (CMC= ∼50 - 100 µM; gray line placed at 100 µM for each plot). nanoDSF experiments (right) reveals that T_m_ decreases upon micelle binding. MST (left) and Nile Red (middle) data were subjected to Bayesian geometric fits with the shading representing 50% and 95% confidence intervals. All experiments were performed in triplicate.

Interestingly, we observe a large change to the *initial fluorescence* that begins at ∼60 µM for OA and DHA, which suggests that a second binding event occurs between FABP7 and either OA or DHA but not SA (Fig. 2). While the *thermophoresis* change occurs over ∼2 orders of magnitude of fatty acid concentration, the *initial fluorescence* change is over a single order of magnitude. The *initial fluorescence* change is too steep for this event to be binding of a second fatty acid ligand.

### FABP7 binds to OA and DHA micelles

After FABP7 is saturated with a singly-bound fatty acid ligands, the free fatty acids in solution may form micelles when the critical micelle concentration (CMC) is reached (24). The exact concentration where micelles spontaneously form can be influenced by the ion concentration, pH and temperature (25). FABPs, including FABP7, can bind to membranes and micelles (14). Indeed, the role of key FABP nuclear localizing residues has been established in part with mutagenesis studies combined with model membranes (8, 26).

We established the CMC of DHA, OA and SA with a lipophilic fluorescent probe Nile red. Free Nile red does not fluoresce in aqueous solution; however, in a lipid rich environment (e.g. when Nile red binds to micelles) the fluorescence intensity increases sharply. The Nile red assays reveal sharp fluorescence intensity changes in the 10^−5^ – 10^−3^ M concentration range.Interestingly, this fluorescence intensity change occurs at similar fatty acid concentrations as the second binding event measured by *initial fluorescence* MST (Fig. 2).

To interpret our experimental data, we utilized a Bayesian geometric programming-based technique to fit the MST and Nile red datasets separately for each ligand (Figs. S3-S5). From these Bayesian fits, we estimate that the OA and DHA CMCs is between 50 and 100 µM. The exact CMC for each ligand is difficult to estimate since the CMCs depend on factors that are difficult to determine experimentally, including the number of fatty acids per micelle, and complex equilibria involving apo and holo FABP7 binding to micelles (Fig. S4). Based on our fitted data of the binding event detected by initial fluorescence and Nile red data establishing the CMC of OA and DHA, we believe that FABP7 is binding to OA and DHA micelles. In contrast to OA and DHA, we do not observe SA form micelles at the fatty acid concentrations probed in the current study (Fig. S2). This absence of SA micelles is consistent with no second binding event being observed between FABP7 and SA by MST.

Based on NMR studies on FABP7s interaction with phospholipid nanodiscs, it has been proposed that *apo* FABP7 (from *Drosophila melanogaster*) has a stronger affinity towards membranes than *holo* FABP7 (15). This suggests that single molecule binding weakens FABP7’s affinity for membranes, and consequently, that single molecule binding and membrane binding are anti-cooperative. We utilized our Bayesian fits of the MST and Nile red data to estimate the equilibria constants of *apo* FABP7 interacting with a micelle (K_D_^P-M^) and *holo* FABP7 interacting with a micelle (K_D_^PL-M^; Fig. S3). We found no correlation between K_D_^P-M^ and K_D_^PL-M^ and can fit a broad range of these values (Fig. S4). From this data alone, we are unable to determine if there is a cooperative, anti-cooperative, or non-cooperative binding between FABP7, single fatty acids, and micelles.

The labeling strategy for MST is *non-specific* and can result in covalent bonding between the fluorophore and any solvent exposed primary amine (N-terminus or K residues) via an N-hydroxy-succinimide ester crosslinking reaction. We labelled a N-terminus 6XHIS-tag of FABP7 with NTA-REDtris, which results in *specific* labelling of the N-terminus far from the main body of the protein. MST performed on *specific* N-terminus labelled FABP7 yields a smaller *initial fluorescence* change compared to *non-specific* labelling (Fig. S1). The difference in initial fluorescence between *specific* and *non-specific* labelling could arise due to changes in the proximity of the fluorophore to the bound micelle.

### FABP7 has a lower melting temperature when associated with micelles, which may indicate micelles uniquely affect protein structure

To determine how micelle-binding affects the structural stability of FABP7, we used nanoscale differential scanning fluorimetry (DSF) to determine melting temperature (T_m_) of FABP7 in the presence of increasing concentrations of OA and DHA. From DSF experiments, two distinct phases are observed (Fig. 2). At equal concentrations of protein and ligand, the protein is stabilized with a small increase in T_m_, which is common upon ligand binding (27). As ligand concentration exceed protein concentration and micelles form there is a decrease in T_m_ for both OA and DHA (Fig. 2). From these data, we observe micelle binding decreases the T_m_ of FABP7, which could be a result of weakened intra-protein interactions. In contrast, SA showed only a small decrease in T_m_ at high fatty acid concentrations, which is likely because SA does not form micelles (Fig. S2).

### FABP7 binds to micelles via residues on helices 2/3 and the βCD turn

To understand how micelles and FABP7 interact, we initiated molecular dynamics simulations with preformed micelles with 50 OA molecules (30 charged and 20 neutral) and apo FABP7 with ∼4.5 microseconds of simulation time. Based on geometric fits on our experimental data, we determined a micelle size of 50 OA molecules to be a reasonable number for our models although the exact size of micelles could vary significantly (Fig. S4).

During the first stages of our simulations, the micelles are not associated with FABP7 and can have vastly different orientations with respect to the protein (Fig. 3a). However, the simulations rapidly converge to structures where the micelles consistently bind to the portal region of FABP7 (Fig 3bc). Specifically, along FABP7’s H3 motif, K21, R30, Q31, N34, and K27 consistently hydrogen bond with the micelle, while interactions are also observed between micelles and the βCD (S55, T56, F57, and K58) and βEF (A75 and D77) turns of FABP7. The residues that most often interact with micelles are near the N-terminus of H3 and the βCD turn (interaction occupancies >75%), while residues in H2 and βEF interact less frequently (occupancies = ∼25-50%) with micelles. In addition to hydrogen-bonding interactions that arise between micelles and FABP7, F27 and F57 intercalate in the hydrophobic micelle interior. Therefore, our simulations indicate that FABP7 – micelle binding is stabilized by both hydrogen-bonding and hydrophobic interactions.

**Figure 3.**
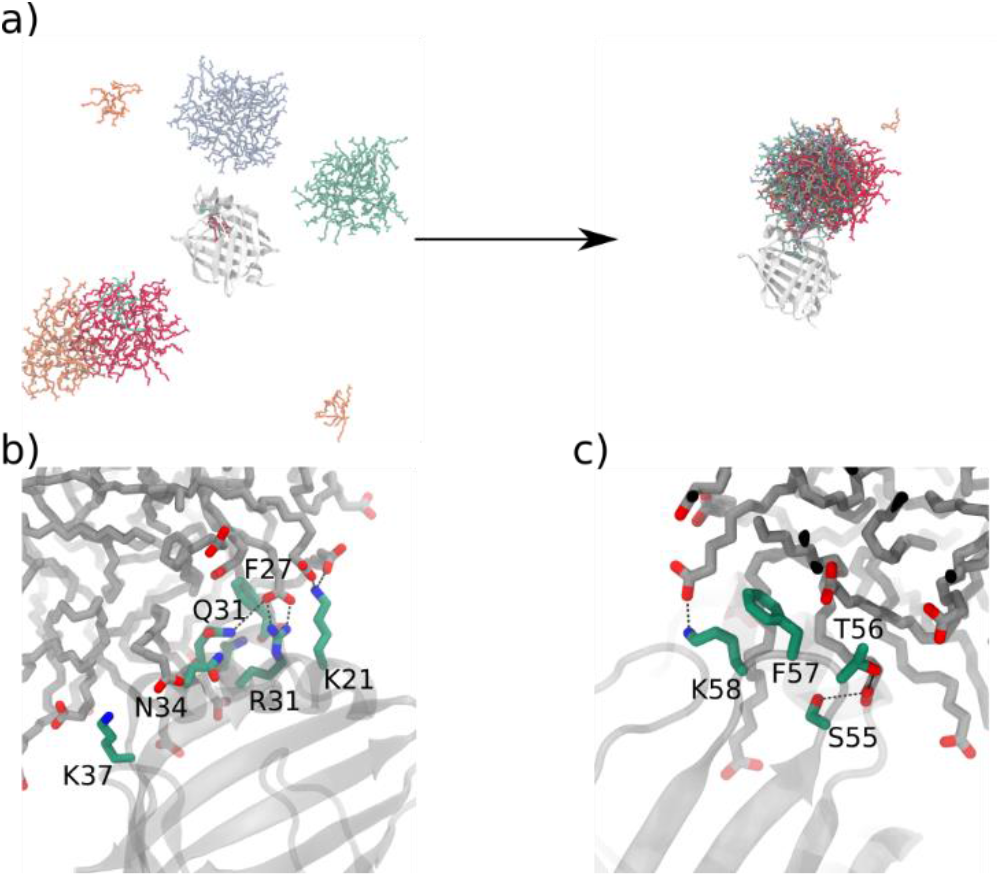
FABP7 interacts with micelles primarily via the βCD loop and H3. a) Several different configurations of unbound OA micelles and FABP7 (after 75 ns of simulation) converge to a single bound conformation (after ∼250 ns of simulation) suggesting that micelles specifically bind to the βCD loop and H3. Hydrogen bonds and non-polar interactions are formed between micelles and b) H3 residues K21, F27, R31, N34, and K37 and c) βCD residues S55, T56, F57 and K58.

### Anomalous NMR titration curves are observed in residues that our simulations predict to bind micelles

We performed 2D ^1^H-^15^N HSQC NMR experiments on ^15^N isotopically labeled FABP7 in the presence of increasing concentrations of OA to detect residue specific chemical shift changes that arise when FABP7 is in the presence of micelles (Figs. 4 and 5). Due to the protein concentration limitations of NMR, we cannot collect data below the CMC of OA and with [FABP7] exceeding [OA], and as a result the minimum concentration of OA and FABP7 is 250 µM and 50 µM, respectively.

**Figure 4.**
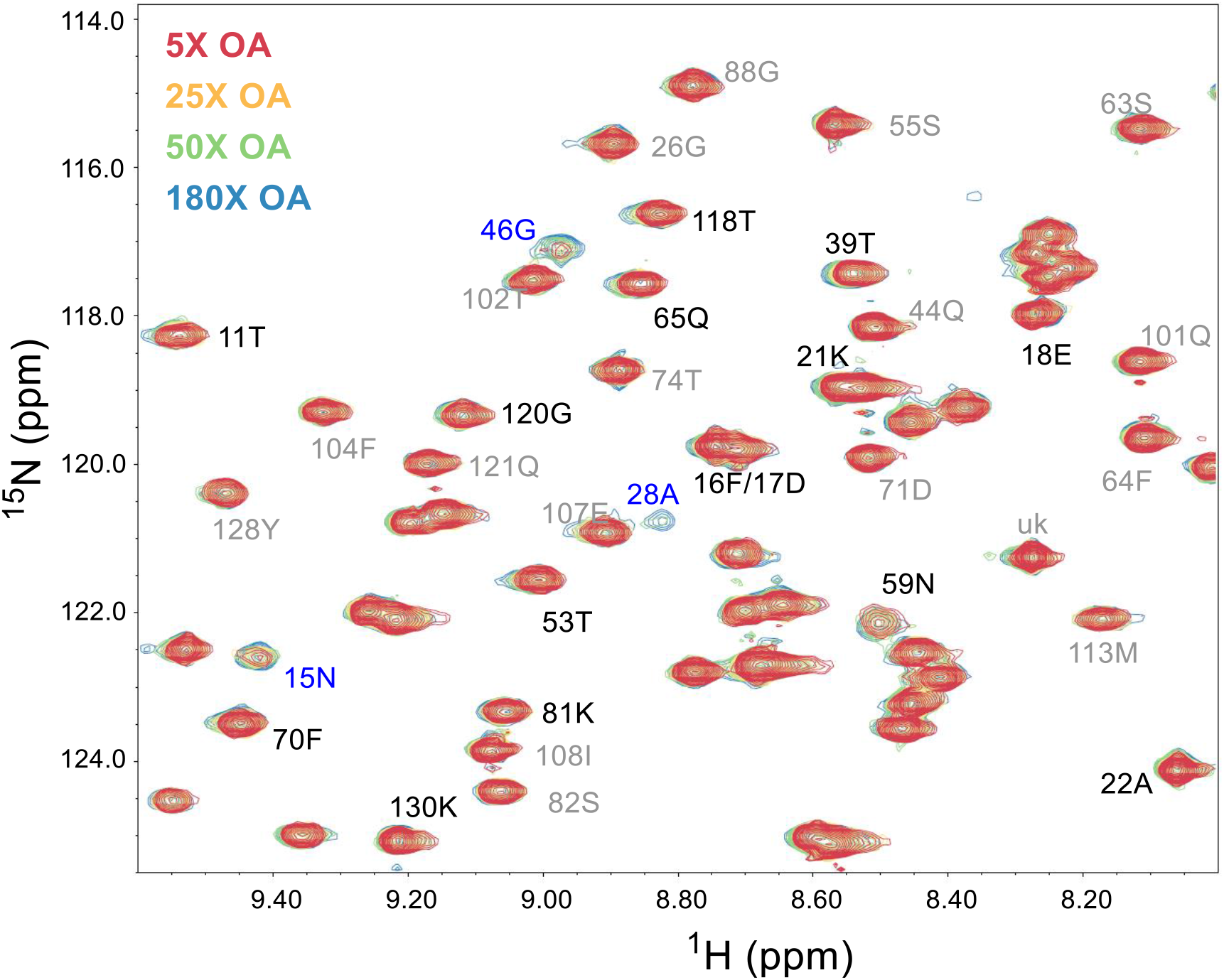
A snapshot of the 2D H N HSQC showing different peak behaviour after OA single molecule saturation. Residue backbone peaks labeled in black show example of residues showing fast exchange CSPs after OA single molecule saturation (maximum peak is followed). Blue highlighted peaks show residues that increase in peak intensity as micelle concentration increases. These spectra are acquired in 50mM phosphate buffer, pH 7.4 with no salt and FABP7 concentrations of 50µM at 25°C.

**Figure 5.**
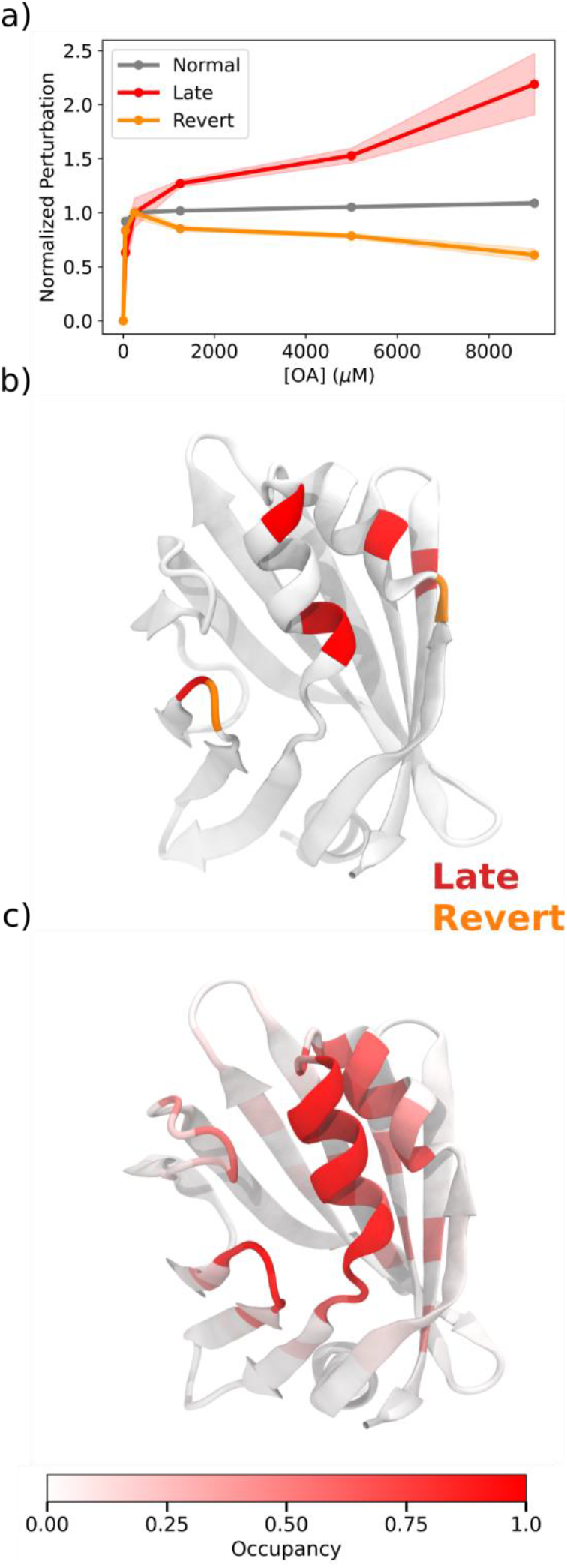
Residues that have significant CSP after binding saturation are consistent with simulation-predicted micelle binding residues. a) Normalized titration curves for 50 µM FABP7 titrated with oleic acid. Curves are normalized to coincide at 250 µM oleic acid (5:1 fatty acid: protein ratio). b) Residues with anomalous titration curves are highlighted on the structure, using the same colour scheme as a). c) Residues that consistently interact with micelles are in H2/H3 and the βCD turn. Occupancy is defined as the fraction of the simulation that an OA molecule resides within 4.5 Å of a residue.

There are many equilibria that contribute to the observed 2D HSQC titration series spectra, including FABP7 binding to single OA ligands or OA micelles. Nevertheless, after [OA] exceeds 250 µM (5:1 [OA]:[FABP7]), CSP changes from single-molecule binding are expected to be minimal since the protein binding pocket is saturated. Certain residues show CSP changes past these concentrations (Figs 4 and 5). Before OA reaches protein saturation (i.e. [OA] = 250 µM and [FABP7] = 50 µM), we observe CSP in slow exchange, which can be attributed to pocket binding of single OA ligands. However, when [OA] exceeds 250 µM, some residues exchange on the fast time scale (Fig. 4, black labels), other residues increase in peak intensity (Fig. 5, blue labels), and certain residues exhibit further CSP changes (Figs. 4 and 5).

The residues with CSP changes after pocket binding saturation are identical or near residues that our simulations suggest bind to micelles (Fig. 5bc). This includes residues on H2 (F16, D17, E18, K21, and A22), H3 (A28), and the βCD turn (N59). We believe that structural data from MD simulations and NMR support a binding model where micelles anchor to the portal region of FABP7, which includes residues on H2 and H3 and the βCD turn.

### FABP7 uses the same residues to bind to micelles and membranes

We probed the interaction of FABP7 and membranes with all-atom and coarse-grained simulations of FABP7 in the presence of model membranes (all-atom: POPC membranes and POPS or POPC membranes for coarse-grained systems). Atomistic simulation permit fine-grain insight into how FABP7 binds to membranes, while coarse-grained simulations allowed us to evaluate how changes to membrane charge and composition affect FABP7-membrane binding.

Our simulations predict that membranes, like micelles, bind to FABP7 through H2, H3, and βCD and βEF turn residues. However, the interactions are less persistent for the membranes for both all-atom and coarse grain systems. Specifically, the interaction between H2/H3 and the membrane is ∼70 % for coarse-grain simulations and ∼65% for all atom simulations. Coarse-grained MD simulations of FABP7 and membranes show more anionic membranes have tighter interactions between FABP7. This is consistent with fluorescence resonance energy transfer (FRET) measurements that show that the rate of transfer of a fluorescently labelled fatty acid between FABP7 and membranes is more rapid with anionic phospholipids present in the membrane (17).

Mutagenesis combined with electron spin resonance experiments have revealed that FABP7 interacts with lysophospholipid micelles through residues in its alpha helices (14). The residues, including D17, E18, K21, N31, and V32, are some of the residues that we predict interact with OA micelles based on NMR and modelling data. Other FABPs also bind to membrane or membrane-mimetics through their H2 and H3. Indeed, for intestinal FABP (FABP2), H2 and H3 are required for membrane binding with helix-less variants losing their affinity for membranes (18, 28). Simulations of ReP1-NCXSQ, another member of the FABP family, reveal that this FABP interacts with membranes with residues in the same regions of the protein (29). Combined with data on both FABP2 and FABP7, this highlights the importance of the portal region for membrane interactions. Our results provide atomistic details on FABP portal region interactions with membranes and thereby provide additional context to a growing body of literature focused on understanding how FABP proteins traffic their cargo throughout the cell. Moreover, given that the nature of the interaction is similar between FABP7 and micelles or membranes, OA or DHA micelles could be a useful model to study how FABPs acquire and deliver ligand cargo to membranes.

### Micelle and membrane binding lead to an open conformation of FABP7

Our atomistic simulations predict that FABP7 binding to micelles or membranes is associated with weaker intra-protein interactions (Fig. S6). For example, residues F57 and K58 primarily form intra-protein contacts with H3 or the βEF turn in the absence of a micelle or membrane. However, when these residues bind to micelles or membranes, the intra-protein interactions are weaker and less persistent and FABP7 adopts an open conformation (Fig. S6). The diminished intra-protein interactions could lead to the lower FABP7 T_m_ that we observe from our nanoDSF data (Fig. 4). Moreover, from our NMR experiments we observe CSP reversion for certain portal region residues when [OA] exceeds [FABP7] by more than 5-fold (Fig. 7, orange). Our simulations predict that the CSP reversion could arise due to FABP7 adopting an open and apo-like conformation when interacting with micelles, which contrasts with the closed conformation that FABP7 adopts when it binds a single OA molecule (Fig. S6).

**Figure 6.**
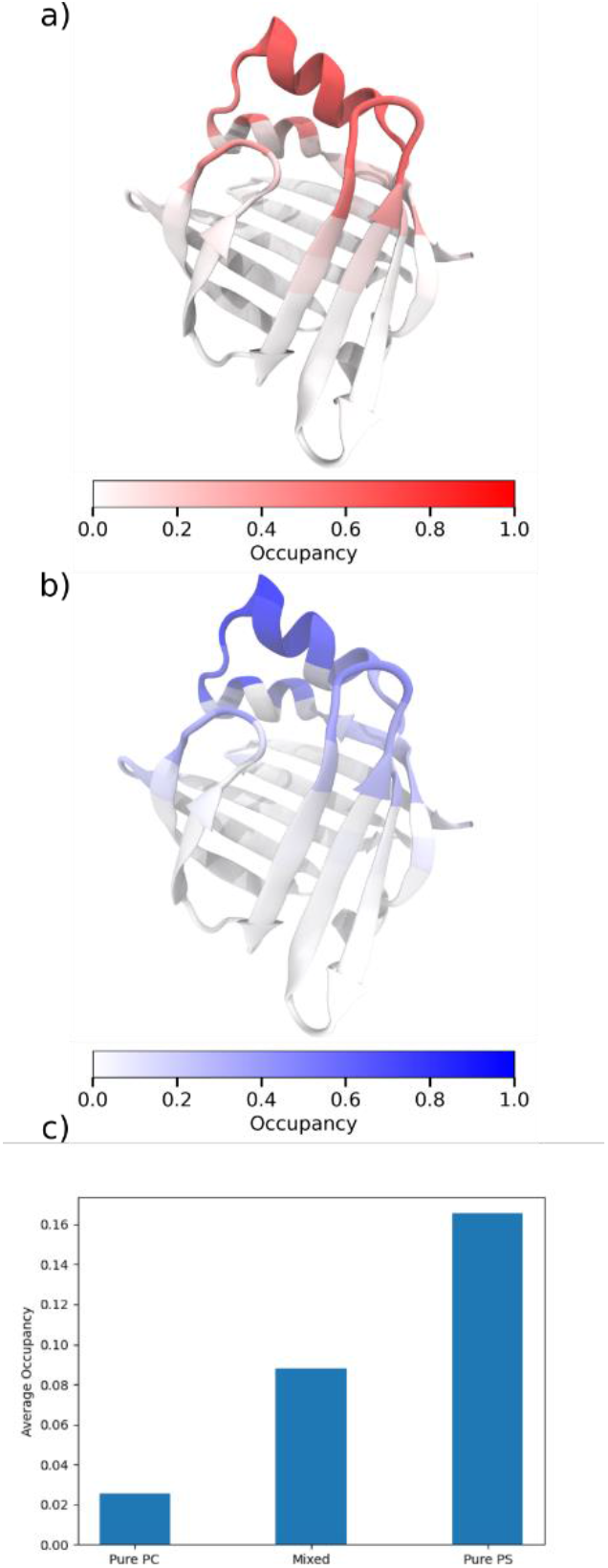
FABP7 interacts with membranes with the same residues it uses to interact with aggregates. FABP7-membrane interactions are primarily through H2-H3 and surrounding portal region residues. a) Average occupancy for each residue mapped onto the structure of FABP7 from atomistic simulations of FABP7 and POPC membranes with 12 DHA molecules. b) Average occupancy for each residue mapped onto the structure of FABP7 from coarse-grained simulations of FABP7 and POPS membranes. Average occupancy for each residue as a function of membrane composition. c) Overall average occupancy from differently charged membranes reveals that FABP7 interacts more frequently with negatively-charged membranes.

**Figure 7.**
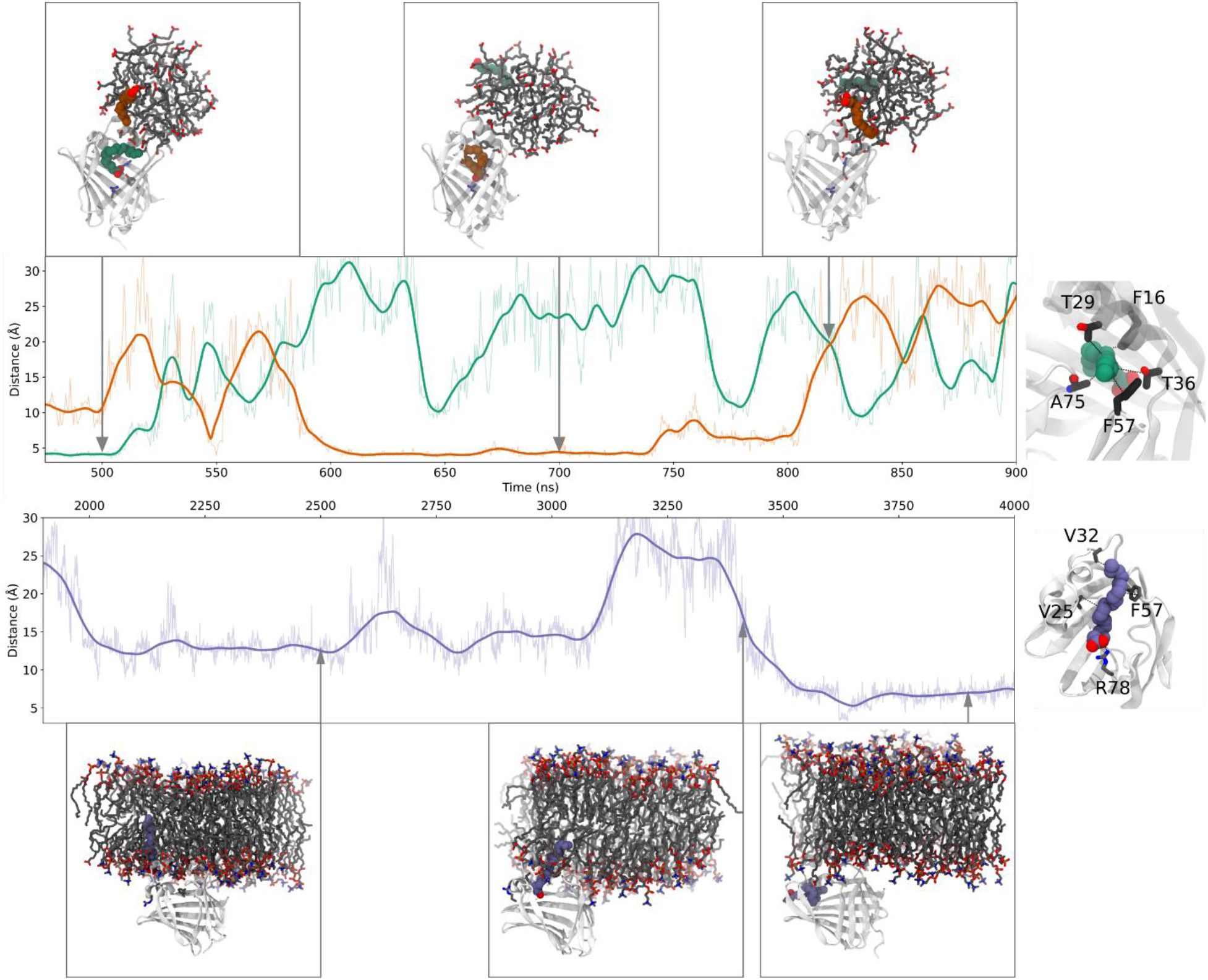
Protein-micelle or membrane association leads to transient pocket binding mediated by portal region residues. Time plot of binding coordinate and two OA molecules. Each line represents the contact distance between a particular OA molecule. The dark line is the smoothed LOWESS curve, while the lighter lines are the raw contact distance. The reaction coordinate is an equally weighted linear combination of the shortest distances between the residues in the right graphic and each OA or DHA molecule.

### Fatty acids enter the FABP7 binding pocket through a space opened by H3 unfolding

From our atomistic simulations, we observe single OA (micelle) or DHA (membrane) molecule reversibly binding to the FABP7 binding pocket from the micelle or membrane. The open conformation assumed by FABP7 in the presence of a membrane or micelle likely permits these ligand binding and unbinding events to occur.

To characterize the binding events, we define a coordinate as a linear combination of contact distances between F16, T29, T36, F57, and A75 and OA molecules for the micelle simulation (DHA and V25, V32, F57, and R78 for membrane simulations). We chose these residues since visual inspection suggested that they chaperone the ligand to the binding pocket. The reaction coordinate is different for membrane simulations due to greater unfolding of H3 and wider gap between βE and βD, which prevents A75 from interacting with the ligand as it proceeds to the binding pocket.

For both membrane and micelle simulations, our data shows that fatty acids enter the FABP7 binding pocket through the space between H3 and the βCD turn. During a micelle simulation, two binding events are observed with two OA molecules that are initially far away from the binding pocket (Fig. 9). Once bound, the ligand adopts a similar conformation that is observed in an FABP7–OA crystal structure (13). For the FABP7-membrane simulation, we observe a single binding event with a DHA molecule entering the FABP7 binding pocket. However, the DHA molecule adopts an alternative conformation where the carboxylate of the fatty acid binds to R78. This alternative pose may be stabilized by the open conformation of FABP7. From all of our membrane sampling data, this is the only instance of a DHA molecule fully disassociating from the membrane and binding to FABP7. Nevertheless, for simulations containing both micelles and membranes, intermediate structures are also detected where the FA is partially inserted into the binding pocket and the tail remains associated with the micelle or membrane.

The fatty acid binding pathways that we identify from our simulations is an example of a collisional transfer of a fatty acid ligand to the binding pocket of FABP. Several FABPs, including FABP7, have been previously shown to bind cargo through a collisional transfer mechanism (18–21). However, the point of entry that ligands use to enter the binding pocket is not known, with studies proposing that ligand enter through the gap between βD and βE, spaces between H3 and the βD and βE turns, and area near the N-terminus (30–34).

Our simulations support a binding mechanism where association of FABP7 with a membrane or membrane-mimetic weakens intra-protein contacts between H3 and βD/βE turns. The weakened intra-protein contacts leads to partial unfolding of H3 (Figs. S6 and S7), which permits fatty acids to enter the FABP7 binding pocket (Fig. 7). The simulations are well-supported by our NMR data that suggests FABP7 associates with micelles through H2/H3 and βCD residues. Furthermore, both nanoDSF and NMR suggest that intra-protein contacts are weakened in the presence of micelles (Figs. 2, 4, and 5), which may be a requirement for cargo uptake. Interestingly, hydrogen-deuterium exchange and chemical exchange saturation transfer (CEST) experiments that show unfolding of H3 is a rate-limiting step for ligand binding to FABP5 (31, 32). While these experiments focused on free ligands in solution binding to an FABP, our data suggests a similar binding pathway exists for ligands binding to FABPs via membranes or membrane-mimetics. Given the common fold that FABPs adopt, we speculate that all FABPs that utilize collisional transfer bind fatty acids through the space between H3 and βCD.

## Conclusion

We used a combined experimental and theoretical approach to examine the nature of the binding interaction between FABP7 and OA or DHA at high fatty acid concentrations. Our biophysical assays indicate that this interaction occurs between OA and DHA micelles binding to FABP7, while our simulations and NMR experiments show this interaction is mediated by FABP7’s portal region. These results show first hand that FABP7 interacts with micelles within an *in vitro* model. Coarse-grained simulations of FABP7 and anionic membranes indicate that the interaction is stronger with increasingly negative charge on the membranes. Our simulations predict that FABP7 can acquire fatty acids from membranes through the space between H3 and the βCD turn. To the best of our knowledge, this is the first study to model the direct transfer of a fatty acid from a membrane/micelle to the binding pocket of an FABP. This knowledge could be used to modulate how FABPs binds to ligands (by mutating residues in H3), which could be used to design specific therapeutics that target FABPs for treatment of inflammation, pain, cancer, or mental disorders.

## Methodology

### Experimental Methods

#### Protein Expression and Preparation

FABP7 wildtype gene sequences were ordered from Life Technologies in the pENTR plasmid that is compatible for GATEWAY cloning. All sequences were cloned into pHGWA expression vector with a tobacco etch virus protease cleavage site inserted between the His tag and protein coding sequence. *Escherichia coli* BL21(DE3) cells were used for the expression of the FABP7-pHGWA. Expression and purification details can be found in the supplementary methods. Final purified FABP7 was dialyzed into 20mM phosphate buffer, pH 7.5, 50mM NaCl.

#### NMR Acquisition

Spectra were collected using a Bruker 600 MHz Avance spectrometer equipped with a 1H, 15N, 13C TXI probe with a single axis z-gradient. Data processing was completed using NMR Pipe and NMR Draw and analyzed with NMR View. All spectra were collected at 25°C and referenced to DSS.

^1^H-^15^N HSQC titration spectra with 50 µM N^15^-FABP7 and OA concentrations of 0, 25, 50, 250, 1250 and 5000 µM were collected for residue specific chemical shift perturbations (CPSs) associated to single OA molecule and micelle binding. Control spectra of FABP7 with the same DMSO concentration showed no difference to apo-HSQC spectra. Assignment of amide backbones was taken from the literature for apo and OA bound FABP (35, 36). Ambiguous and overlapping peaks were excluded from this study.

Weighted chemical shift perturbations between apo and fatty acid bound spectra were calculated as follows:

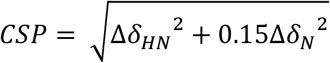

Where Δ*δ*_*HN*_ and Δ*δ*_*N*_ is the chemical shift difference between apo and holo spectra for the proton and nitrogen dimension, respectively. For residues that have multiple peaks in the apo spectra, the most intense peak was used for the CSP calculation.

#### Ligand preparation

OA and DHA was purchased from Sigma, and SA from Larodan. Stock solutions and 1:1 serial dilution were all prepared in DMSO. Fatty acid-DMSO were mixed with assay solutions to a maximum 2% DMSO content. All dilution series were completed in low-bind Eppendorf tubes.

#### Microscale Thermophoresis

Microscale thermophoresis (MST) measurements were carried out using Monolith NT.115Pico instrument. Measurements were carried out in MST-buffer (20 mM phosphate, 120 mM NaCl, at pH 7.4, in 0.005% Tween-20) using premium capillaries (Nanotemper). Proteins were labeled with RED-tris-NHS dye following supplier protocol. Measurements were taken with instrument set to high MST power, and 20-30% LED power.

A solution of 20 nM FABP7-NHS-RED-Tris labeled was combined with fatty acid dilution series. Samples were incubated for 30 minutes at measurement temperature before loading samples into MST capillaries.

Analysis of MST traces was completed using MO.Affinity Analysis v2.3 (Nanotemper) (37). Relative florescence (F_norm_) is determined using the same defined time intervals for the initial and thermo-diffusion florescence. All measurements are completed in triplicate.

*Nile Red Assay*. Nile Red critical micelle determination experiments were carried out in the Cytation 5 Plate Reader (BioTek) in black 96-well plates (Greiner). The ligand dilution series of concentrated OA, DHA, and SA were performed similar to the method described above, except Nile Red was diluted in the DMSO to have a final concentration of 200 mM in the assay tubes. A solution of PBS/Tween 20 (20 mM phosphate, 120 mM NaCl, at pH 7.4, in 0.005% Tween-20) was combined with ligand/DMSO/Nile red in low-bind tubes, vortexed and pulse spun. Samples were incubated for 30 minutes at measurement temperature prior to transfer to the plate and measured.

#### Nano-DSF Measurements

Nano-DSF measurements were carried out using the TYCHO instrument (Nanotemper). Similar to the MST experiments, protein, with and without ligand was loaded into micro-capillaries by dipping the end into the solution and placing the capillaries into the analysis tray and closed into the tray. The TYCHO measures intrinsic fluorescence (330 and 350 nm) with increasing temperature as the protein melts. The change in fluorescence at these two wavelengths and the ratio of their values can be used to determine the protein’s melt temp. Unlabelled FABP7 wild-type protein was diluted to 20 μM in PBS (20 mM phosphate, 120 mM NaCl, at pH 7.4). The protein was combined with the fatty acid ligands in low-bind tubes in the same manner as with the MST experiments. Samples were incubated for 30 minutes at measurement temperature prior to transfer to the capillaries. All measurements were performed in triplicate.

### Computational Methods

#### FABP7 Starting Structure

The X-ray crystal structure of FABP7 bound to antinociceptive SBF1-26 (PDB ID: 5ura)(38) was used as a starting structure for our models containing FABP7. For simulations of apo FABP7 without ligands, micelles, or membranes, the system was neutralized with 4 Na^+^ ions, and solvated with a 10 Å octahedral water box, which results in ∼5000 waters with a side length of ∼61 angstroms. For all atomistic models, the bound inhibitor was removed from the binding pocket and hydrogen atoms were assigned using H++.(39)

The CHARMM-GUI (40) web server was used to prepare inputs for simulations of FABP7 with micelles and membranes.

#### Micelle Simulation Setup

To generate an equilibrated micelle structure, initially 50 oleic acid molecules (30 charged, and 20 neutral) were placed in a water box with 9786 water molecules and 30 Na^+^ using the Multicomponent Assembler tool from CHARMM-GUI. The system was subjected to minimization, heating, and equilibration (see below for more details) and simulated for 500 ns.

#### Micelle and FABP7 Simulation Setup

The equilibrated micelle and FABP7 protein were placed in a water box with 9326 water molecules and 34 Na^+^ counterions using the Multicomponent Assembler tool from CHARMM-GUI.

#### Atomistic Membrane and FABP7 Simulation Setup

Using the Membrane Builder tool from CHARMM-GUI, a system containing FABP7 and a POPC membrane with interspersed DHA was constructed. First, the membrane was aligned along the Z-axis, while the FABP7 protein was translated 50 Å along the Z-axis. A 40 Å band of water was placed above and below the membrane, while the X and Y side lengths were assigned to be 65 Å. The outer leaflet of the membrane contained 60 POPC molecules and 12 DHA molecules, while the inner leaflet contained 65 POPC molecules and 0 DHA molecules.

#### Atomistic Simulation Details

All atomistic molecular dynamics simulations were run using the OpenMM python library (41). The simulations were performed using Langevin dynamics with a timestep of 2 fs (1 fs for heating and equilibration runs) and a collision frequency of 1 ps^-1^.

Electrostatic interactions were treated with particle mesh ewald (PME) and non-bonded interactions were cutoff using a force-switch cutoff method (10 - 12 Å). Bonded terms involving hydrogen were constrained during production runs. Minimization steps were performed with the limited-memory Broyden-Fletcher-Goldfarb-Shanno algorithm. The pressure was maintained at ∼1 bar with a Monte Carlo Barostat (isotropic for micelle simulations and anisotropic for membrane simulations). For production runs, data was analyzed after 250 ns of equilibration every 50 ps.

#### Minimization, Heating, and Equilibration Protocol

Each system was subjected to three rounds of minimization (5000 steps) with 1) the protein restrained (k = 41840 kJ mol nm^-2^); 2) only the protein sidechains restrained ((k = 10460 kJ mol nm^-2^); and 3) unrestrained. The systems were then heated gradually from 10 K to 310 K over 550 ps with a 10460 kJ mol nm^-2^ restraint placed on the protein. The restraint was then released over 550 ps.

#### Forcefield Details

Apo and holo simulations without the presence of a micelle or membrane were assigned parameters from the ff14SB forcefield (42). The OA ligand was optimized with B3LYP-6-31G(d) using Gaussian 16 (Revision B.01) (43). Subsequently, the ligand was assigned GAFF (44) parameters with AM1-BCC charges using the Antechamber utility from Amber Tools. For simulations containing a micelle or membrane, the CHARMM36m forcefield (45) was used for the protein and lipid parameters were obtained from CHARMM-GUI (40). For all atomistic simulations, explicit water molecules were assigned TIP3P parameters.

#### Coarse-grained Simulation Details

Systems containing FABP7 and membranes were set up using CHARMM-GUI (40). All coarse-grained systems contained FABP7, ∼6000 water molecules, and 0.15 M NaCl and a membrane that was either completely POPC (256 molecules), POPS (256 molecules), or ∼80% POPC (205 molecules) and ∼20% POPS (51 molecules). Each system was assigned parameters from the Martini 2.0 forcefield and minimized and equilibrated using the default scheme provided by CHARMM-GUI. Briefly, this entails 5000 steps of steepest descent minimization, and several rounds of equilibration where the timestep was adjusted from 5 ps to 20 ps. For all coarse-grained simulations, temperature and pressure were controlled at 303.15 K and 1 atm, respectively, using Berendsen thermostat and barostat. Each system was simulated across 7 runs for a total of ∼4 µs of sampling time per system.

#### Analysis

All analysis was performed with either the CPPTRAJ (46) module from Amber Tools or the mdtraj python library (47).

## Supporting information

supplementary methods, figures, tables, and references

