## supplementary methods, figures, tables, and references for "FABP7 Binds to Fatty Acid Micelles: Implications for Lipid Transport"

Plasmid and protein sequence

Protein Expression and Purification

To express, an overnight inoculant was subcultured into LB media at 37°C until cells reached an OD600 of 0.6-0.8. Expression was then induced with 1 mM IPTG and the sample was incubated at 18 °C for 18-20 hours. Cell pellets were resuspended in lysis buffer (25 mM imidazole, 3mM DTT, 50 mM sodium phosphate, 250 mM NaCl, pH 7.8) and lysed with five minutes of sonication. Soluble fraction was separated from cellular debris with centrifugation (40,000g, 30 minutes 4ºC). His-labelled FABP7 was then immobilized with Ni-NTA IMAC column (AKTA system, GE) and eluted in high imidazole solution (250 mM imidazole, 50 mM sodium phosphate, 250 mM NaCl, pH 7.8). The protein was incubated overnight with TEV at 4 °C and Histag and TEV were purified out with Ni-NTA beads (GE). Because FABP7 binds to hydrophobic molecules, the recombinant protein must be delipidated to remove endogenous bacterial lipids. This was performed by two successive incubations of FABP7 with Lipidex beads (Perkin Elmer) for one hour each at 37 °C. Purified protein was then dialyzed into a final solution (20 mM sodium phosphate, 120 mM NaCl, pH 7.4).

### Supplementary Figures

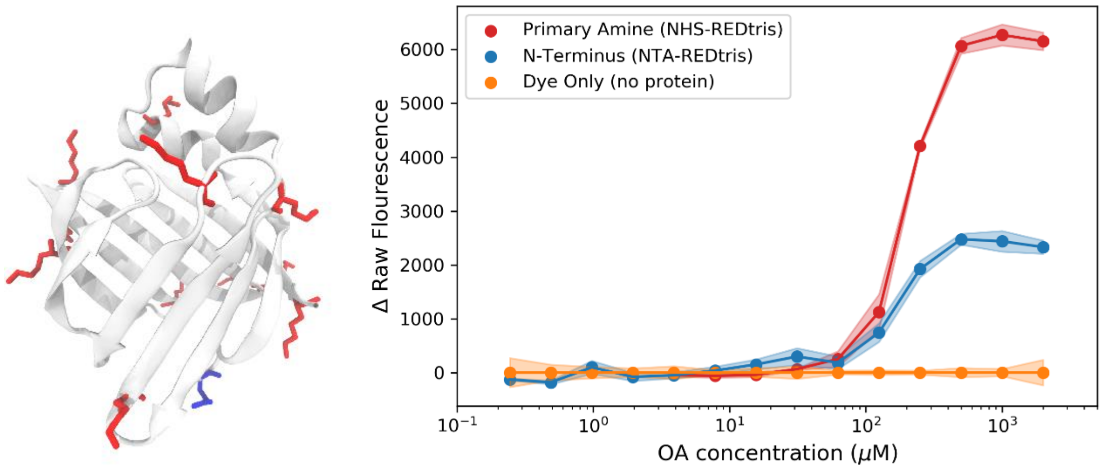


Figure S1. Specific N-terminus RED labelling strategy results in a lower change in initial fluorescence indicating fewer interactions to the N-terminus. All lysine residues available for non-specific labeling of wtFABP7 with NHS-REDtris label are shown in red. Blue shows where the N-terminus of FABP7 will be labeled with NTA-REDtris label. Difference in the initial fluorescence response for different labeling strategies and no protein dye control (n=3). All measurements performed at 37°C.

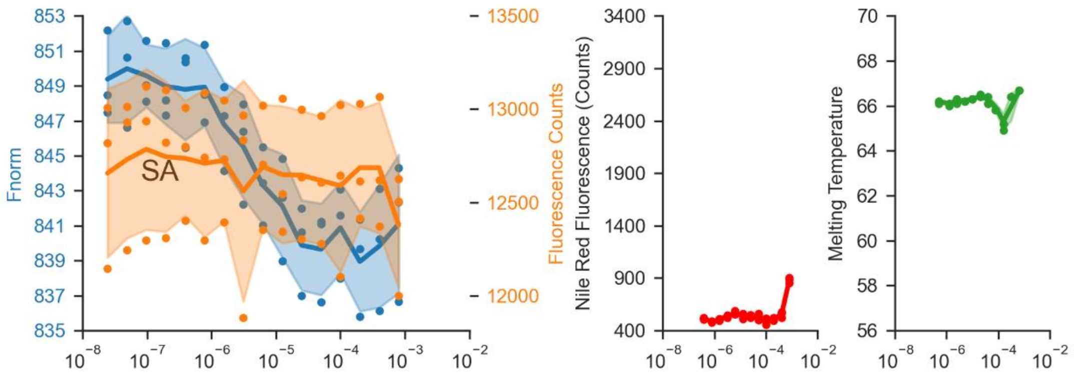


Figure S2. Dilution series experiments probing SA binding to FABP7 reveals no second binding event since SA does not form micelles at the concentrations examined. Thermophoresis (left, blue) and initial fluorescence (left, orange) plots show significant thermophoresis changes (indicative of pocket binding) but little changes to initial fluorescence . Nile red assay (middle, red) determination of the critical micelle concentration reveals that SA does not form micelles at the concentrations examined. nanoDSF experiments (right) reveals that $T_m$ is relatively constant with increasing [SA]. All experiments were performed in triplicate.

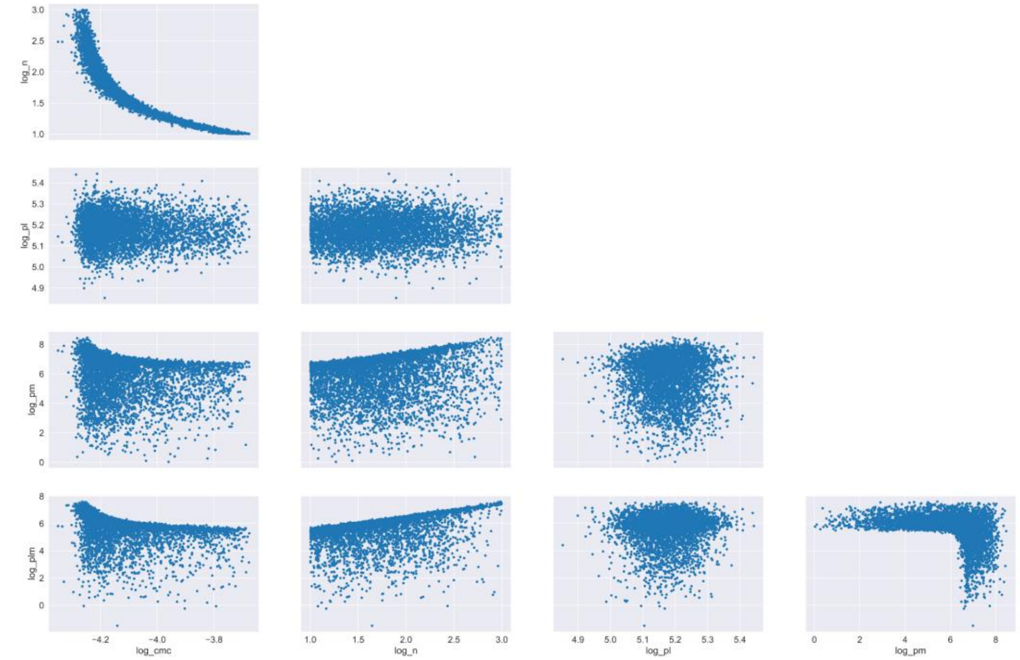


Figure S3. There are strong correlations between some of the fit parameters.

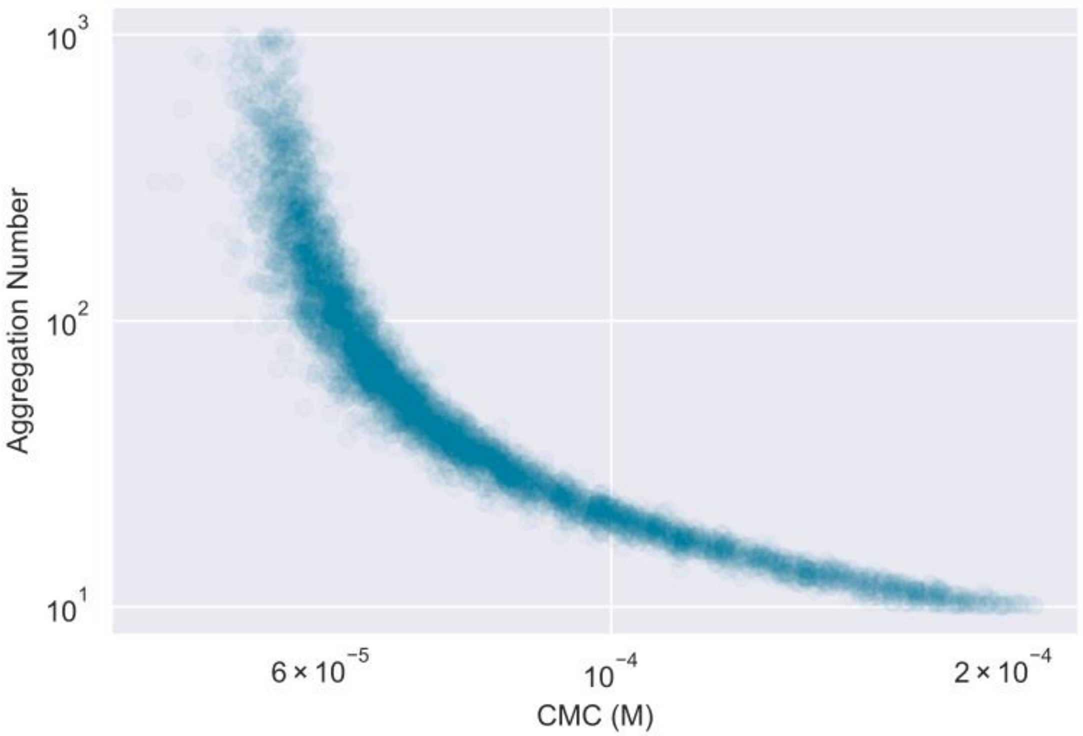


Figure S4. It is difficult to determine the exact CMC and aggregation number from our data.

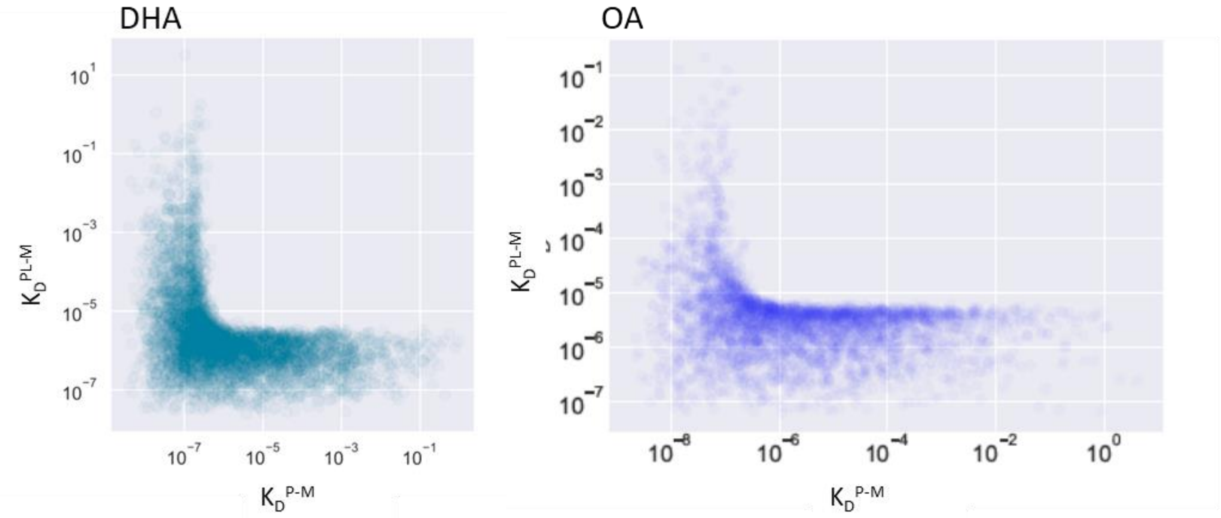


Figure S5. Our thermophoresis and TRIC measurements cannot determine if there is cooperativity or anti-cooperativity between ligand binding and micelle binding.

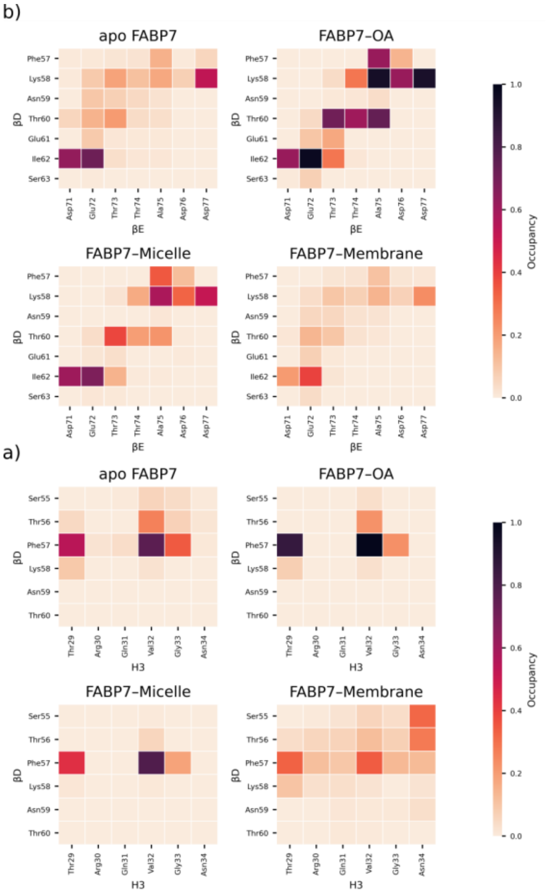


Figure S6. Contact map graph showing contact distance and occupancy between βCD and βEF loops for apo, OA-holo and OA-aggregate simulations.

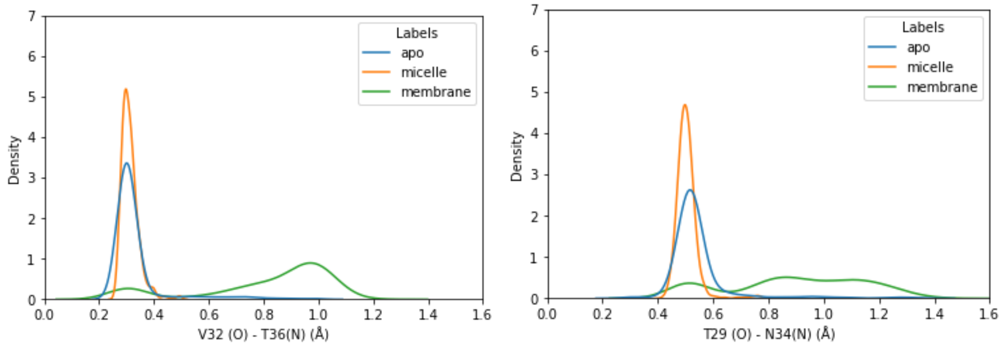


Figure S7. Kernel density plots of the distances between V32 and T36 (left) and T29 and N34 (right) for apo-FABP7, FABP7-micelle, and FAPB7-membrane simulations
